# ABC-HuMi: the Atlas of Biosynthetic Gene Clusters in the Human Microbiome

**DOI:** 10.1101/2023.09.18.558305

**Authors:** Pascal Hirsch, Azat Tagirdzhanov, Aleksandra Kushnareva, Ilia Olkhovskii, Simon Graf, Georges P. Schmartz, Julian Hegemann, Kenan Bozhüyük, Müller Rolf, Andreas Keller, Alexey Gurevich

## Abstract

The human microbiome has emerged as a rich source of diverse and bioactive natural products, harboring immense potential for therapeutic applications. To facilitate systematic exploration and analysis of its biosynthetic landscape, we present ABC-HuMi: the Atlas of Biosynthetic Gene Clusters (BGCs) in the Human Microbiome. ABC-HuMi integrates data from major human microbiome sequence databases and provides an expansive repository of BGCs compared to the limited coverage offered by existing resources. Employing state-of-the-art BGC prediction and analysis tools, our database ensures accurate annotation and enhanced prediction capabilities. ABC-HuMi empowers researchers with advanced browsing, filtering, and search functionality, enabling efficient exploration of the resource. At present, ABC-HuMi boasts a catalog of 19,218 representative BGCs derived from the human gut, oral, skin, respiratory and urogenital systems. By capturing the intricate biosynthetic potential across diverse human body sites, our database fosters profound insights into the molecular repertoire encoded within the human microbiome and offers a comprehensive resource for the discovery and characterization of novel bioactive compounds. The database is freely accessible at https://www.ccb.uni-saarland.de/abc_humi/.

**GRAPHICAL ABSTRACT:** 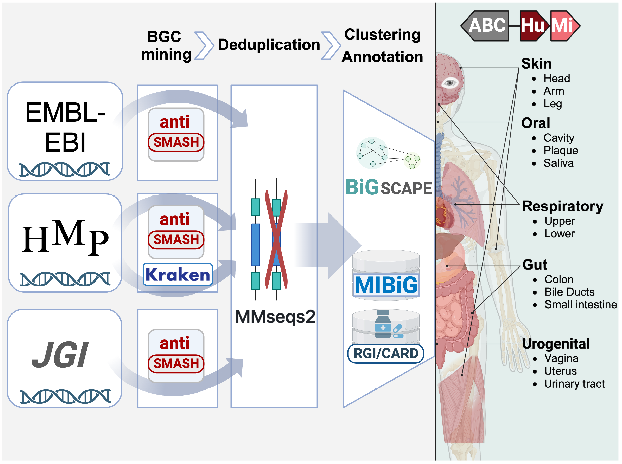

## INTRODUCTION

Bioactive compounds produced by the human microbiome play key roles in host-microbe and microbe-microbe interactions and might greatly affect the health state of an individual (1, 2, 3, 4). These compounds are often encoded in biosynthetic gene clusters (BGCs) harbored by multitudes of bacterial species populating the human body. The identification and thorough analysis of BGCs provide insights into the biosynthetic capabilities of the microbiome and ultimately lead to the discovery of pharmaceutically relevant compounds (5, 6, 7, 8, 9, 10). Community initiatives such as the Human Microbiome Project (11, 12) accumulated vast volumes of sequencing data whose biosynthetic potential still has to be explored.

The genome mining software paved the way for the creation of databases of BGCs computationally predicted from voluminous genomics datasets (13, 14, 15, 16, 17). Some of these databases, such as antiSMASH-DB (13), IMG-ABC (14), and BiG-FAM (15), collect BGCs from a wide variety of sources, while others target specific environments, such as the human gut (16) or ocean microbiome (17). However, there is still no dedicated human microbiome BGC database going beyond the most studied human gut environment and spanning multiple body sites. Furthermore, most of the existing databases were created with already outdated software and do not benefit from the latest advances in genome mining techniques.

To address these limitations, we developed ABC-HuMi, the Atlas of BGCs in the Human Microbiome. Our resource accumulates data from three major human microbiome sequence databases and represents BGCs originating from five human body sites and systems. We employed the most advanced BGC mining and analysis software to populate the database and created a user-friendly interactive platform for exploring the collected BGCs and their associated functionalities.

## DATA RETRIEVAL AND PROCESSING

### Data sources

ABC-HuMi integrates data obtained from the Genomic Catalogue of Earth’s Microbiomes (JGI GEM (18)), EMBL-EBI MGnify catalogs (19), and the Human Microbiome Project (HMP, (20)). The EMBL-EBI MGnify Unified Human Gastrointestinal Genome v2.0.1 and Human Oral v1.0 catalogs contain more than 290,000 isolates and MAGs from the human microbiome clustered into 4,744 and 452 species representatives, respectively. For further processing, we retrieved the representative genomes (https://www.ebi.ac.uk/metagenomics/browse/genomes). Selecting only the representative genomes as the input is a compromise between performance and sensitivity. This allowed us to keep the total number of BGCs small, but our experiments show that we still cover the vast majority of the metabolomic diversity present in the skipped genomes (75% at the gene cluster family (GCF) level and 98% at the gene cluster clan (GCC) level) (Supplementary Figure S1).

JGI GEM contains more than 52,000 MAGs from a wide range of environmental and host-associated microbiomes. We retrieved all human microbiome-associated MAGs, excluding the ones associated with the human gut, already covered by the EBI data (https://portal.nersc.gov/GEM/genomes/).

HMP contains metagenomes from more than 3,808 samples associated with different parts of the human body. From this dataset, we selected all metagenomes associated with respiratory, skin, and urogenital samples (https://portal.hmpdacc.org/). We also included 1,692 HMP reference genomes available at NCBI (BioProject PRJNA28331) ranging in quality from complete genomes to draft contig assemblies. The genomes without associated body site metadata or related to the diseased patients were not considered.

### Processing pipeline

As an initial processing step, we harmonized metadata obtained from the source databases and unified the body site categories. For EMBL-EBI MGnify catalogs, the body site was retrieved from the metadata of associated NCBI BioSamples. Metagenome assemblies were taxonomically annotated using Kraken2 (v2.1.2; command-line arguments (cla): --confidence 0.1) and its PlusPF database (v20220908) (21).

The retrieved genomes were processed with antiSMASH (v7.0.0; cla: --genefinding-tool=prodigal-m --asf) (22). A BGC was classified as fragmented if it is located on a contig edge and complete otherwise. All BGCs with the same body site and taxonomy were grouped and MMseqs2 (v14.7) (23) was used to compare BGCs inside each group. We then merged BGCs with a minimum sequence identity of 0.95 into a single database entry with combined metadata. Finally, BGCs were clustered with BiG-SCAPE (v1.1.5; cla: --mibig) (24) using Pfam (v35.0) (25). We predicted antibiotic resistant genes located within all BGCs at three reliability levels (Perfect/Strict/Loose) using the Resistance Gene Identifier (RGI) software (v6.0.3; cla: -- include loose) of the Comprehensive Antibiotic Resistance Database (CARD) database (v3.2.8) (26). GC content of BGCs and the corresponding full genome sequences was computed with BioPython (v1.5.3) (27).

### Overview of the collected data

The resulting database comprises 19,218 BGCs grouped into 8,989 GCFs and 294 GCCs. Figure 1 shows the composition of the database with regard to the associated body site and product types of the BGCs.

**Figure 1.**
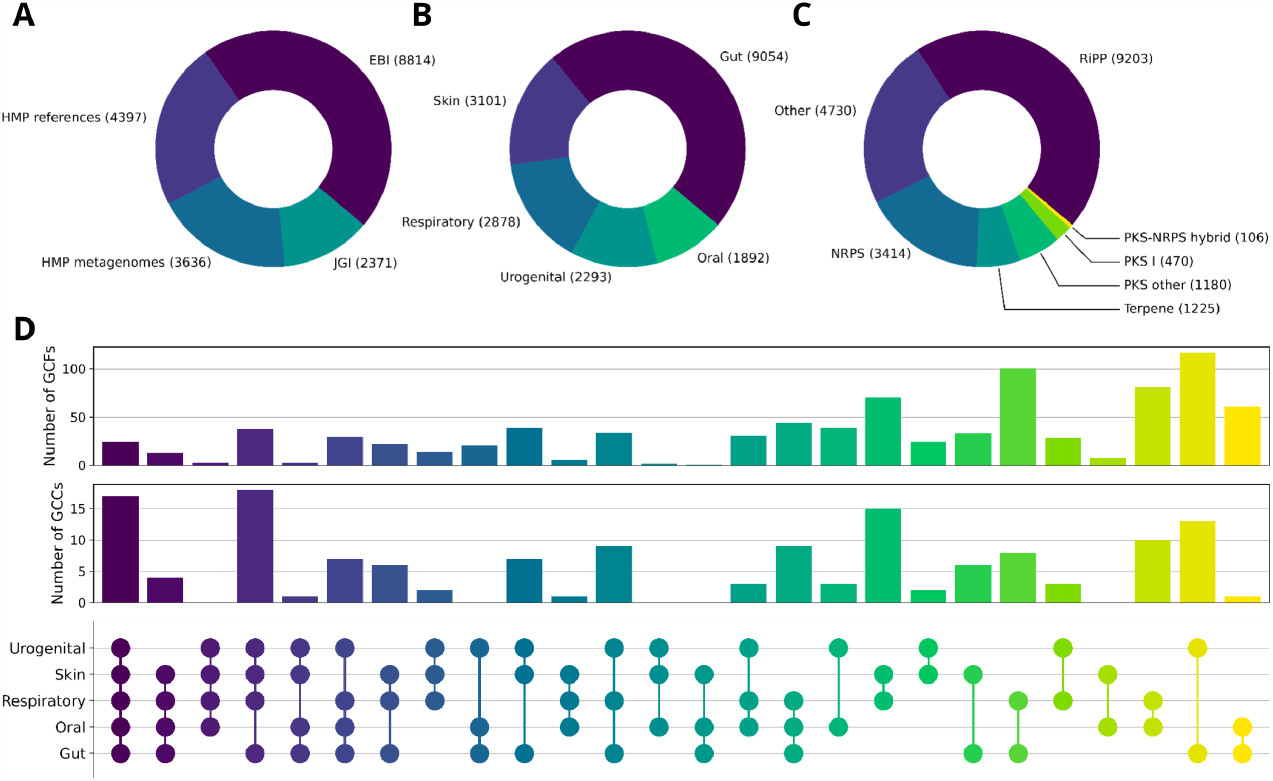
Overview of the ABC-HuMi content. The distributions of biosynthetic gene clusters (BGCs) by (**A**) data source, (**B**) human body site, and (**C**) product type. (**D**) The number of gene cluster families (GCFs) and clans (GCCs) that span across at least two different body sites.

### Web server implementation

For the implementation of the ABC-HuMi web server, we set up a Django Python web framework (https://djangoproject.com/) and a PostgreSQL database (https://www.postgresql.org/) in docker containers (https://www.docker.com/) with the help of a Cookiecutter template (https://cookiecutter.readthedocs.io/). For the search job query, we are using the task queue manager Celery (http://docs.celeryproject.org) together with the in-memory data structure store Redis (https://redis.io/). The search jobs are handled with BLAST+ (28) using the Biopython wrapper (https://biopython.org/docs/dev/api/Bio.Blast.Applications.html) and cblaster (29) using a Snakemake pipeline (30). The database browsing table is using DataTables (https://datatables.net/) and the Cytoscape network visualization (31) is using cytoscape.js (https://js.cytoscape.org/). The front end of the website also uses Bootstraps (https://getbootstrap.com/) and Font Awesome (https://fontawesome.com/) for design purposes and jQuery (https://jquery.com/) as a utility library.

## DATABASE FUNCTIONALITY

### Versatile search options

ABC-HuMi provides the BLAST+ (28) search for identifying homologs of user-provided sequences in the database, for example, antibiotic resistance genes (Figure 2A, B). The function handles both nucleotide and protein sequences and can also search translated nucleotide sequences of the database BGCs using a protein query (the tBLASTn mode). Since the single gene search is rarely informative for identifying homologous gene clusters (32), we employ cblaster (29) for screening a custom BGC against the entire database (Supplementary Figure S2-3).

**Figure 2.**
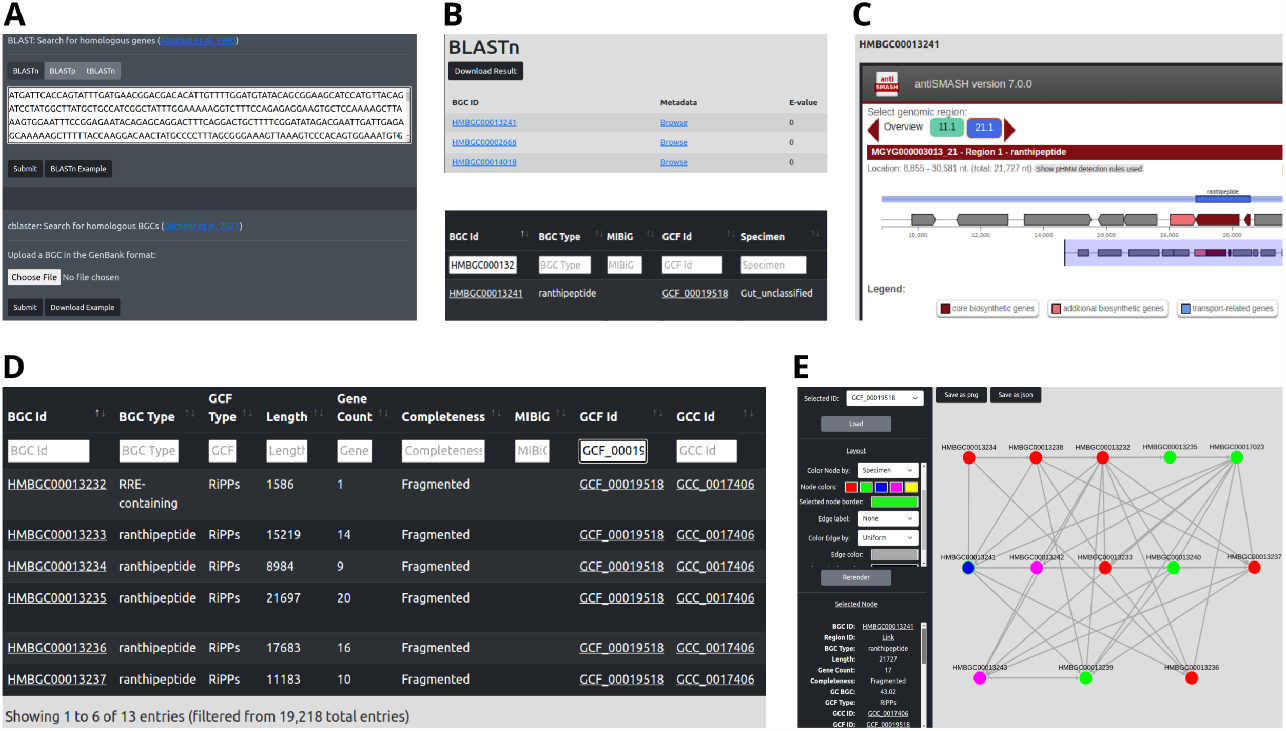
ABC-HuMi search, browse, and visualization functionality. The BLAST search (**A**) identifies biosynthetic gene clusters (BGCs) harboring homologs of the user-provided sequence. The search result page (**B**, top) enables browsing of the BGC metadata (**B**, bottom) and the corresponding embedded antiSMASH reports (**C**). The metadata table links to the gene cluster family (GCF) or clan (GCC) of a BGC, thus allowing one to browse all similar clusters (**D**). GCC and GCF networks can also be visualized via interactive Cytoscape plots (**E**). Here all nodes of a GCF are titled with the BGC identifiers and colored according to the corresponding human body site: the selected blue node is from the gut, green and pink nodes are from the oral microbiome and red ones are from the respiratory system.

### Interactive tables and visualizations

All BGCs in ABC-HuMi are enriched with detailed metadata that augments antiSMASH-predicted properties with the source human microbiome sequence database and body site, the taxonomy of the producing organism, and the links to similar BGCs within our database and in MIBiG (33). The information is stored in interactive tables that allow filtering by metadata and seamless switching between BGCs and related gene cluster families (GCFs) and clans (GCCs) (Figure 2B, D). Each BGC is complemented with the fully-embedded antiSMASH report; GCF and GCC networks are visualized with Cytoscape (Figure 2C, E).

### Application examples

#### Exploring individual BGCs

To explore the hidden bioactive potential of GCFs spanning across multiple human body sites, we employed the versatile ABC-HuMi search functionality. First, we used metadata filtration and Cytoscape visualizations to identify GCFs containing experimentally-validated MIBiG BGCs whose direct neighbors originated from distinct body sites. We further selected two out of 11 hits since the corresponding MIBiG BGCs are known for producing compounds with antimicrobial properties: bacteriocins gassericin T (BGC0000619) and gassericin E (BGC0001388). Both compounds were originally discovered in *Lactobacillus gasseri* (34), one of the main *Lactobacillus* species identified in the vaginal, gastrointestinal, and oral microbiomes (35). The cblaster search of BGC0000619 and BGC0001388 revealed four BGCs in ABC-HuMi with identifiers HMBGC00006519-22 (Supplementary Figures S2-3). The body sites associated with these BGCs are gut, oral and urogenital system, thus perfectly matching the common localization of the known gassericin producer. Notably, three of the ABC-HuMi BGCs were predicted from *L. paragasseri*, never reported as a gassericin E producer before. Given the resemblance between these two bacterial species, such production is plausible and the ABC-HuMi BGCs might harbor a new variant of the bacteriocin. Though, this computational hypothesis requires experimental validation.

#### Large-scale comparison

To demonstrate the ABC-HuMi applicability to large-scale analysis, we explored BGCs in a recently published dataset of gut microbiota samples derived from patients with Parkinson’s disease (26). First, we applied our data processing pipeline to the binned and unbinned metagenomic data from (26) split into three cohorts of individuals according to the study metadata. BGCs identified with antiSMASH in each group were further used as queries for bulk comparisons against the entire ABC-HuMi database (Supplementary Figure S4). The high ratio of unmatched BGCs at the GCF level (32-43%) strikes the enormous diversity of the human gut microbiome which is not fully covered by the MGnify catalogs of representative genomes. The small difference in distributions of matched BGCs in healthy and diseased cohorts agrees with the original study results, where no significant difference in the microbiome diversity of cohorts was discovered (26). At the same time, the ABC-HuMi-based analysis highlights BGCs shared by multiple body sites (11-14%). This additional layer of information was hidden in the original study focused exclusively on the human gut microbiome but might represent interest for further exploration.

### COMPARISON TO EXISTING RESOURCES

With over 2500 entries, MIBiG is by far the leading community resource for collecting experimentally-verified BGCs (33, 36). However, such BGCs are scarce. A more complete — though less accurate — view on the biosynthetic potential of environmental microbiomes relies on genome mining (5), which naturally leads to the emergence of resources storing computationally-predicted BGCs, such as our database. These resources visibly differ in data focus, processing techniques, and functionality provided to the end users. We compare the ABC-HuMi content and features to sBGC-hm, the only human microbiome-specific BGC database to date, and three state-of-the-art general-purpose *in silico* BGC databases (Table 1).

**Table 1.** Comparison of *in silico* BGC databases. *# HM-related BGCs (GCFs)* stands for the total number of BGCs (GCFs) related to the human microbiome.

| Category | ABC-HuMi | sBGC-hm | BiG-FAM | antiSMASH-DB | IMG-ABC |
| --- | --- | --- | --- | --- | --- |
| *General information* |  |  |  |  |  |
| Data focus | Human microbiome | Human gut microbiome | Microbes in general | Microbes in general | Microbes in general |
| Genome types | isolates, MAGs, metagenomes | isolates, MAGs | isolates, MAGs | isolates | isolates, MAGs |
| Latest update | 2023 (this work) | 2023 (16) | 2020 (15) | 2020 (13) | 2019 (14) |
| *Data processing and availability* |  |  |  |  |  |
| BGC calling | antiSMASH v7 (22) | antiSMASH v6 (37) | antiSMASH v5 (38) | antiSMASH v5 (38) | antiSMASH v5 (38) |
| BGC clustering | BiG-SCAPE (24) | BiG-SCAPE (24) | BiG-SLICE (39) | ✗ | ✗ |
| # HM-related BGCs (GCFs) | 14,821 (7346) | 36,583 (8004) | >4791 (>811)<sup>a</sup> | unknown | unknown |
| Bulk data download | ✓ (GenBank) | ✗ | ✓ (custom) | ✗ | ✗ |
| *Features* |  |  |  |  |  |
| Metadata search | ✓ | ✓ | ✓ | ✓ | ✓ |
| Sequence search | ✓ (BLAST+ (28)) | ✓ (BLAST+ (28)) | ✗ | ✗ | ✗ |
| Cluster search | ✓ (cblaster (29)) | ✗ | ✓ (BiG-SLICE (39)) | ✓ (ClusterBlast (40)<sup>b</sup>) | ✗ |
| BGC detailed view | ✓ (antiSMASH) | ✗ | ✓ (custom<sup>c</sup>) | ✓ (antiSMASH) | ✓ (antiSMASH<sup>d</sup>) |
| Extra features | Cytoscape visualization of GCF/GCC networks with export to PNG and JSON | Matrices of gene co-occurrence in the HMP data<br>Visualizations of differences in core gene coverage across HMP samples | Word cloud of Pfam features for each BGC | Cluster search based on NPRS/PKS module queries<br>Taxonomy tree-based browsing | Pfam-based search with ClusterScout (41)<br>Search by chemical attributes (for experimentally-verified products) |

<sup>a</sup>The exact numbers are known for human gut MAGs only.<sup>b</sup>Indirectly available via the antiSMASH pipeline and web interface.<sup>c</sup>BGC annotation into genes and Pfam domains.<sup>d</sup>Unavailable for part of the database BGCs.

A direct comparison of the total number of BGCs across the databases could be misleading because of the drastic differences in the focus of the resources, the sourced microbiomes, and the types of underlying genome sequences. AntiSMASH-DB includes only BGCs predicted from isolate genomes, other databases also consider MAGs and ABC-HuMi further relies on unbinned metagenome assemblies as the mining source. The latter increases the number of less reliable and fragmented BGC predictions but enables deeper exploration of the hidden potential of the microbiome. ABC-HuMi keeps the information about the genome type behind each predicted BGC, so users can easily adjust the data selection to their research demands.

While all five databases rely on antiSMASH for mining BGCs, the software version varies from v5 (BiG-FAM, antiSMASH-DB and IMG-ABC) to v6 (sBGC-hm) to the latest v7 (ABC-HuMi), which might greatly affect the accuracy and completeness of the predictions (22, 37, 38). ABC-HuMi, sBGC-hm, and BiG-FAM group identified BGCs into gene cluster families (GCFs) and clans (GCCs). The first two resources utilize BiG-SCAPE (24) while the enormous size of BiG-FAM requires the use of more resource-efficient but less accurate BiG-SLICE (39). The resulting BGCs could be downloaded in bulk only from the ABC-HuMi and BiG-FAM websites, but the further use of the BiG-FAM data might be complicated due to the non-standard data format.

All considered databases provide comprehensive filtering by metadata but the search for a custom sequence or BGC is available only in three of them (ABC-HuMi, BiG-FAM, and sBGC-hm) and only ABC-HuMi supports all search types. Though, antiSMASH-DB and IMG-ABC offer advanced data lookups based on a user-defined set of Pfam domains from a predefined list. AntiSMASH-DB can also be indirectly queried for a custom BGC via the ClusterBlast algorithm embedded into the antiSMASH pipeline (40). The visualization facilities vary greatly among the databases but the most informative detailed view of each BGC (the antiSMASH report) is provided only in ABC-HuMi and antiSMASH-DB.

## CONCLUSION

The unprecedented biosynthetic potential of the human microbiome remains underexplored. In response, we unveil ABC-HuMi, a repository of computationally predicted biosynthetic gene clusters (BGCs) from major human body sites and systems. Despite its compact size, the database represents a huge biosynthetic diversity of the human microbiome and provides a convenient framework for its exploration. By looking at BGCs whose GC content is substantially different from the average genome GC, users might detect possible horizontal BGC transfer events. By inspecting antibiotic resistance genes co-located with BGCs, researchers might identify the producers of promising bioactive compounds. By exploring the distribution of similar BGCs across body sites, one can shed light on the general-purpose or niche-specific function of the underlying BGCs.

The streamlined nature of ABC-HuMi facilitates easy and fast analysis queries, even for computationally demanding applications. The database empowers users with the ability to search not only for custom nucleotide and protein sequences but also entire BGCs and enhances the results with interactive browsing and visualization functionality. Unlike most existing BGC databases, the entire ABC-HuMi can be downloaded locally and used for bulk comparison and prioritization of novel BGCs in large-scale studies. We thus believe the created resource will be beneficial for the diverse needs of the rapidly growing community. We plan to maintain and further expand ABC-HuMi in both functionality and coverage of the human microbiome biosynthetic diversity.

## CODE AND DATA AVAILABILITY

The code used to collect and process the data can be found at https://github.com/gurevichlab/abc_humi. The resulting annotated BGC sequences in the GenBank format and all metadata are available from the database website. The resource is freely accessible at https://www.ccb.uni-saarland.de/abc_humi/.

## SUPPLEMENTARY DATA

Supplementary Data are available online.

## ACKNOWLEDGEMENTS

The graphical abstract was created with BioRender.com. We thank our colleagues at HIPS for testing the ABC-HuMi database functionality and suggesting new features.

