## Supplementary Data for "ABC-HuMi: the Atlas of Biosynthetic Gene Clusters in the Human Microbiome"

[1]Chair for Clinical Bioinformatics, Saarland University, Saarbrücken 66123, Germany, [2]Helmholtz Institute for Pharmaceutical Research Saarland (HIPS), Helmholtz Centre for Infection Research, Saarbrücken 66123, Germany, [3]Saarbrücken Graduate School of Computer Science, Saarland University, Saarbrücken 66123, Germany, [4]Department of Computer Science, Saarland University, Saarbrücken 66123, Germany, [5]Department of Pharmacy, Saarland University, Saarbrücken 66123, Germany

---

*To whom correspondence should be addressed. (R.M.), (A.K.), (A.G.) †These authors contributed equally to this work

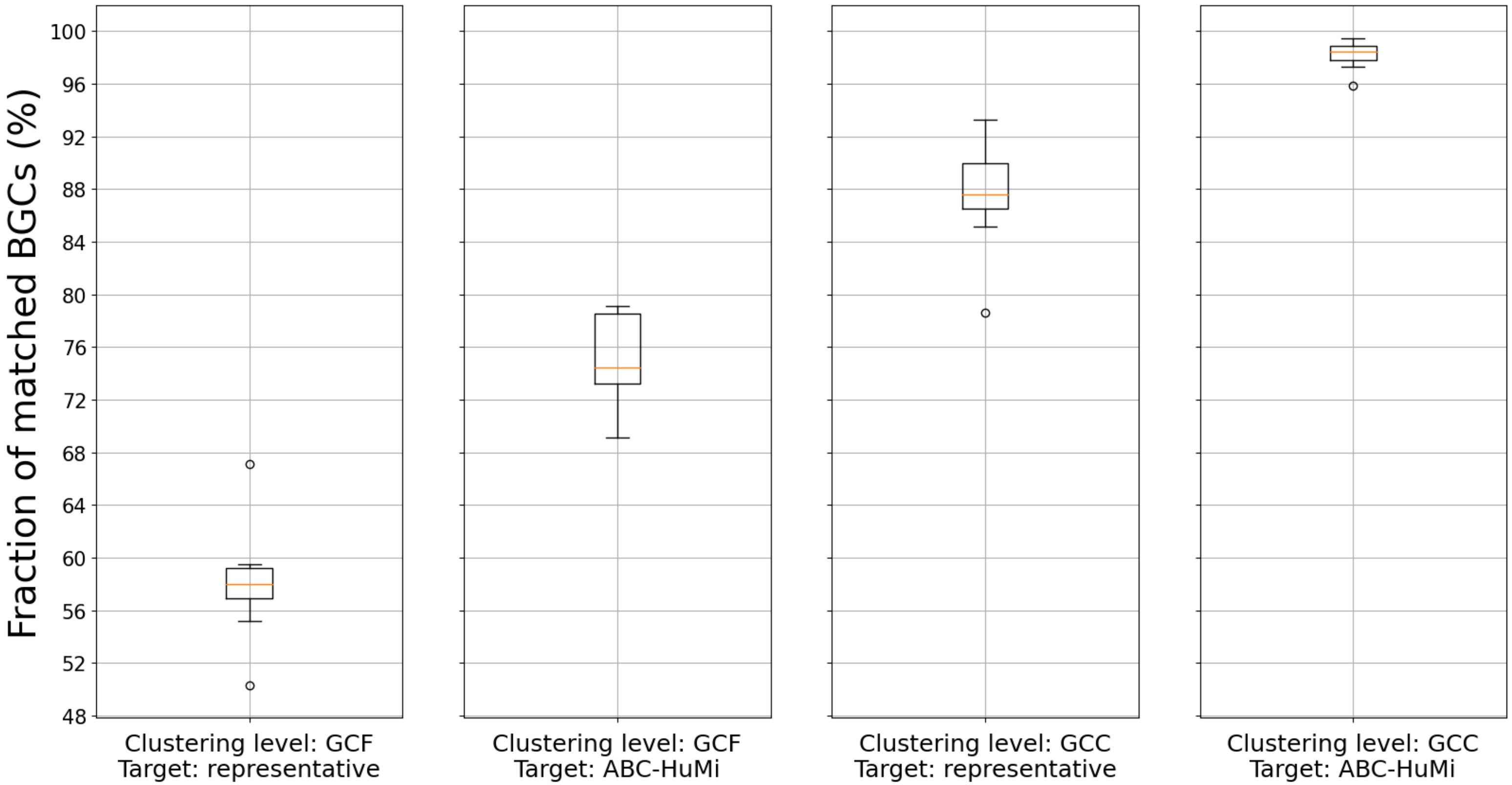


**Supplementary Figure S1**. To investigate how well representative genomes represent BGCs contained in genomes from the EMBL-EBI MGnify [1] Human Gut v2.0.1 catalogue, we randomly sampled 10 groups of 100 non-representative genomes and analyzed them with our pipeline. The BGCs contained in these groups were clustered by BiG-SCAPE [2] together with BGCs from (1) the respective representatives and (2) the ABC-HuMi database. In this figure, we show distributions of a fraction of non-representative BGCs that were matched with target BGCs for different clustering levels.

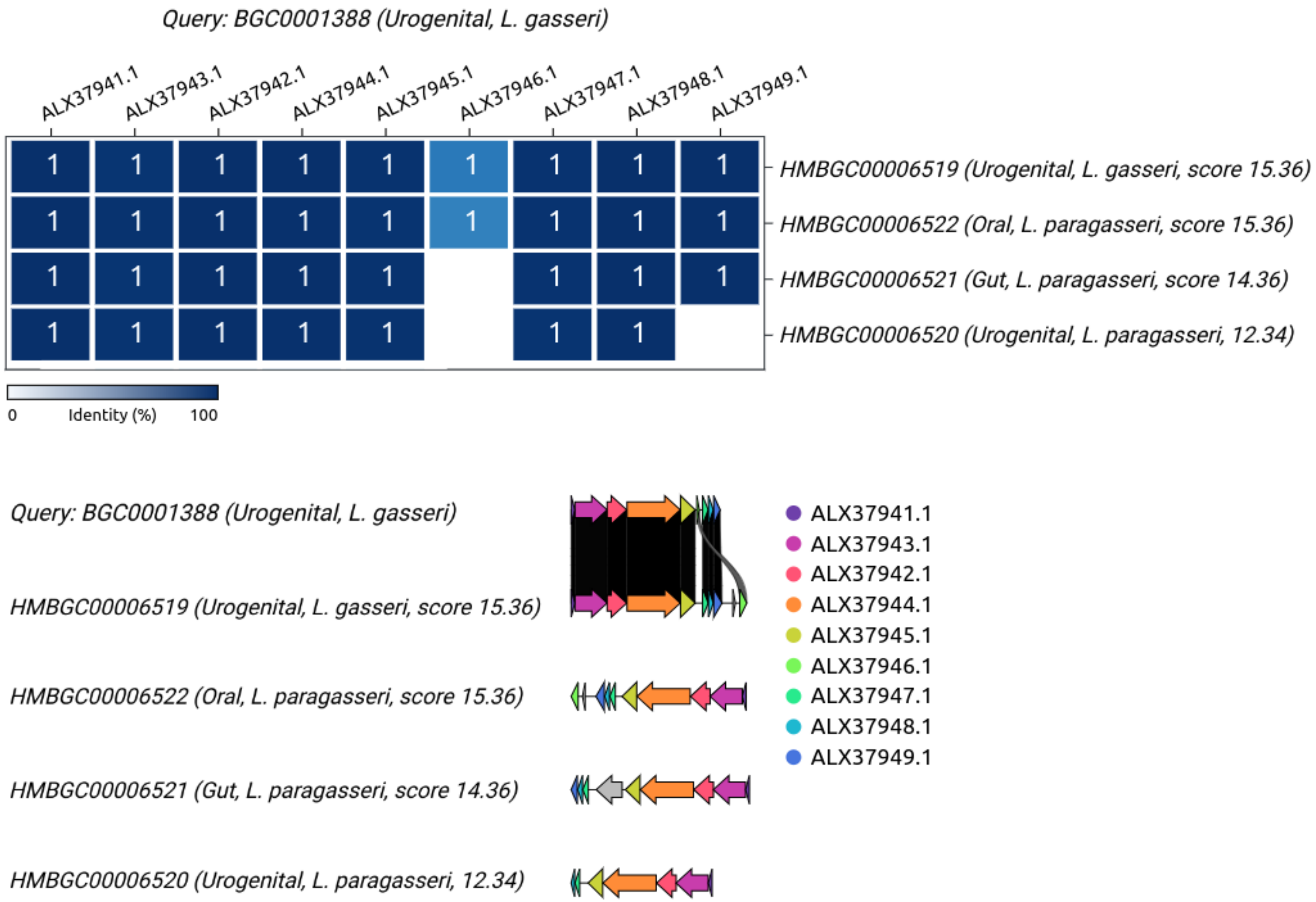


**Supplementary Figure S2**. Results of the cblaster [3] (top) and clinker [4] (bottom) searches of the gassericin E biosynthetic gene cluster (MIBiG ID: BGC0001388) [5]. Only hits with pairwise gene sequence identity >95% are shown.

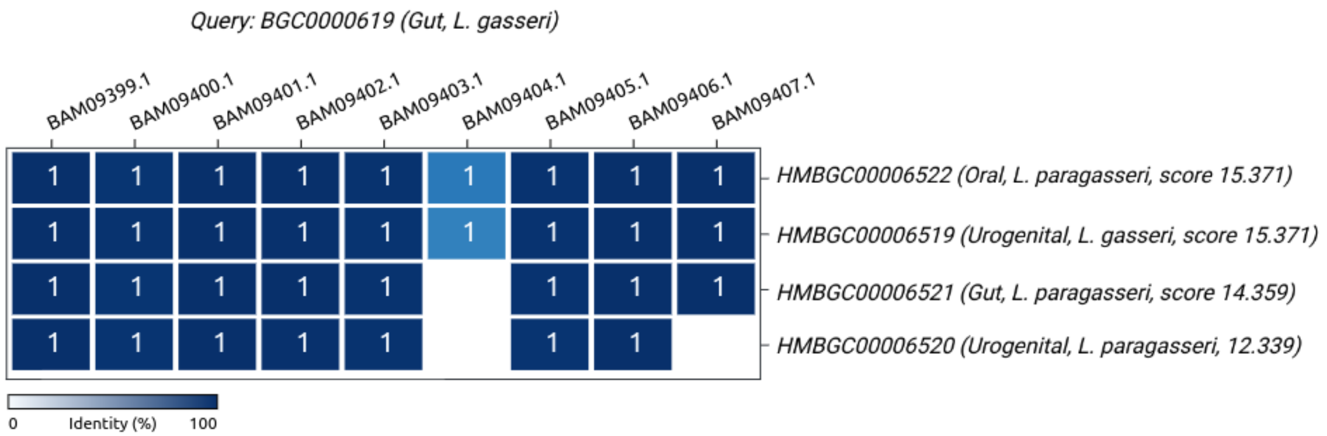


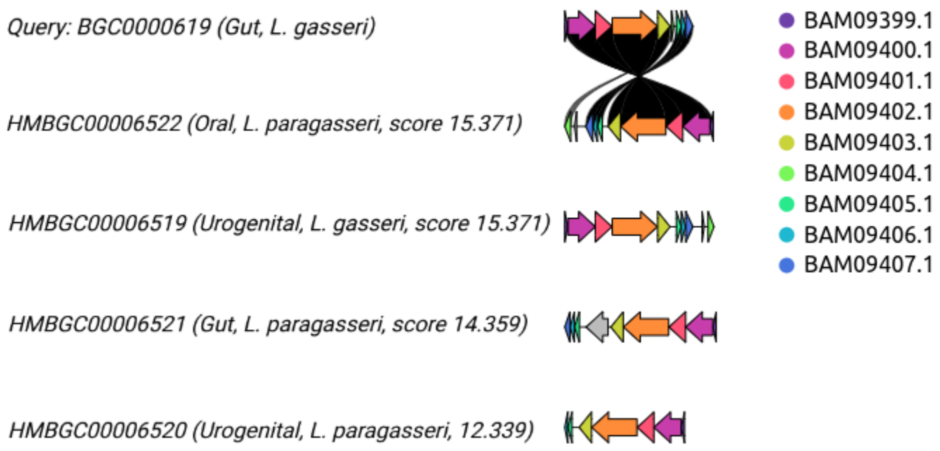


**Supplementary Figure S3**. Results of the cblaster (top) and clinker (bottom) searches of the gassericin T biosynthetic gene cluster (MIBiG ID: BGC0000619). Only hits with pairwise gene sequence identity >95% are shown.

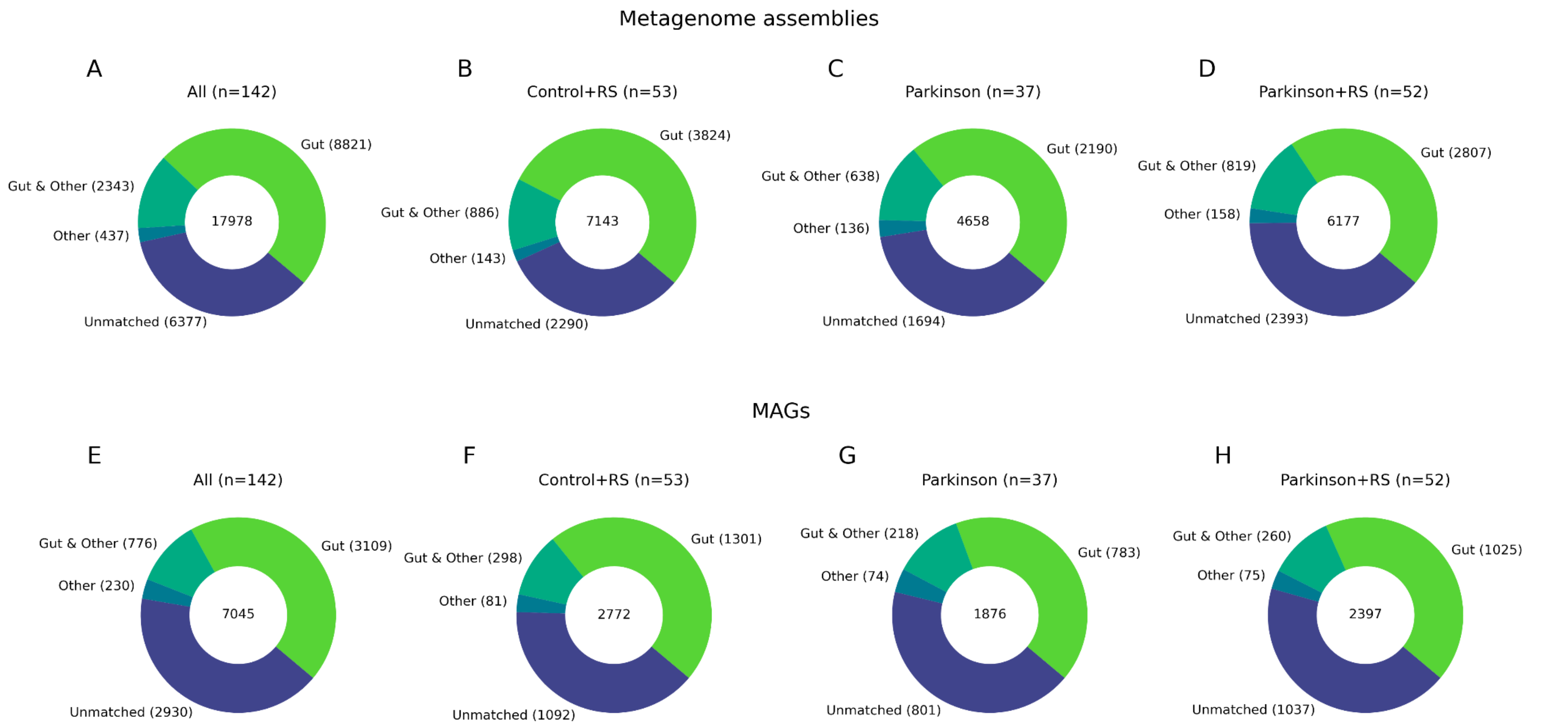


**Supplementary Figure S4**. The fraction of BGCs in the RESISTA-PD study [6] clustered with the ABC-HuMi BGCs at the gene cluster family (GCF) level (the BiG-SCAPE default threshold of at most 0.3 similarity). The panels show numbers of BGCs identified in samples from all (**A, E**), healthy (**B, F**) and two diseased (**C, D, G, H**) cohorts of individuals, categorized by the body site linked to ABC-HuMi BGCs (Gut, all Other sites, and both). BGCs unique to the RESISTA-PD study are classified as unmatched. The total numbers of the study BGCs in each cohort are given in the circle centers. The results are computed for BGCs derived from metagenome assemblies (**A-D**) and MAGs (**E-H**).
